# Neuroendocrine differentiation (ND) in sensitivity of neuroendocrine tumor (NET) cells to ONC201/TIC10 cancer therapeutic

**DOI:** 10.1101/2024.08.28.610183

**Authors:** Elizabeth Ding, Maximillian Pinho-Schwermann, Shengliang Zhang, Connor Purcell, Wafik S. El-Deiry

## Abstract

Prostate cancer (PCa) neuroendocrine tumor (NET)-like cells with low or absent androgen receptor (AR) signaling cause hormone therapy resistance and poor prognosis. Small cell lung carcinoma (SCLC), a high-grade NET, presents with metastasis early and has poor survival. ONC201/TIC10 is a first-in-class cancer therapeutic with clinical activity in diffuse gliomas and neuroendocrine tumors. We hypothesized that markers of neuroendocrine differentiation, activation of the integrated stress response (ISR) and the TRAIL pathway, as well as the expression of ClpP, contribute to neuroendocrine tumor cell death and sensitivity to ONC201. We show that PCa and SCLC cell lines (N=6) are sensitive to ONC201, regardless of the extent of neuroendocrine differentiation. Endogenous levels of some NET markers (CgA, FoxO1, ENO2, PGP9.5, SOX2) are present in a spectrum in PCa and SCLC cell lines. Overexpression of neural transcription factor BRN2 in DU145 PCa cells does not increase expression of NET differentiation markers FoxO1, ENO2, PGP9.5, and CgA at 48 hours. However, the transient BRN2 overexpression showed slight decreases in some NET markers on the spectrum while maintaining sensitivity of PCa cells to ONC201 before any phenotypic change related to NET differentiation. Our results show that ONC201 has preclinical activity against PCa including those without NET markers or in PCa cells with transient overexpression of neural transcription factor BRN2. Our results have relevance to activity of ONC201 in PCa where most castrate-resistant androgen-independent cancers are not therapy resistant due to NET differentiation. Importantly, NET differentiation does not promote resistance to ONC201 supporting further clinical investigations across the spectrum of PCa.

## Introduction

Prostate cancer (PCa) is the second most common malignancy and the fifth leading cause of cancer death among men. There are an estimated 1.4 million new cases and 375,000 annual deaths due to the disease worldwide [1]. Advancement of PCa to metastatic disease significantly relies on the androgen receptor (AR). Androgen deprivation therapies (ADT) remain a mainstay for treating metastatic PCa patients by ablating AR signaling [2]. ADT show efficacy in around 90% of patients through a decrease in prostate-specific antigen (PSA) levels [3]. While localized prostate cancer can be removed or treated successfully, treatments for advanced metastatic prostate cancer remain challenging [4].

After a mean of 2-3 years, prostate cancer resurfaces as castration-resistant prostate cancer (CRPC) [3]. Reactivation of the AR occurs despite low serum androgen, resulting in disease progression and metastasis [2]. For patients with advanced metastatic castration-resistant prostate cancer (mCRPC), prognosis is poor and mean survival time is around 25.6 months from diagnosis [5]. Mainstay treatments for mCRPC have expanded to include second-generation AR pathway inhibitors (API), like enzalutamide (ENZ) and abiraterone (ABI), and further suppress AR signaling. While it has shown to increase overall patient survival [6], API resistance invariably follows second-line treatment, and complete treatment is rare ([2], [7]).

API-resistant prostate adenocarcinomas are a significant clinical challenge due to scarcity of third-line therapeutics and its potential in progressing into lethal neuroendocrine prostate cancer (NEPC) via neuroendocrine differentiation (NED) [2]. Neural transcription factor BRN2 has been shown to be a master regulator of NED and NEPC development *in vitro* through NE marker expression and CRPC progression [8]. Previous studies have shown that AR directly represses BRN2, which affects BRN2 regulation of SOX2 [2], an additional NEPC-related transcription factor [9].

Only 0.5-2% of NEPC arise *de novo*, while around 25% of advanced PCa are NEPC from trans-differentiation of prostate adenocarcinoma, CRPC cells, or cancer stem cells due to therapy resistance [10]. Through lineage alterations, PCa cells develop neuroendocrine cell characteristics, including expression of neuroendocrine markers like chromogranin A (CgA) [11], PGP9.5/UCHL1 [12], synaptophysin (syp), BRN2, FOXA1, enolase 2 (Enol-2), and more [10]. While patients show decreased PSA levels due to AR signaling loss [13], NEPC patients show poor prognosis with high metastatic burden, lack of therapeutics, and a <20% rate of 5 year survival [6]. Thus, there is a great need for effective therapies for high-risk neuroendocrine prostate cancer.

ONC201 is a first-in-class imipridone that was initially discovered as a TNF-Related Apoptosis Inducing Ligand (TRAIL)-inducing compound TIC10, but later shown to initiate the integrated stress response (ISR) [14] through ATF4/CHOP [15], inactivates Akt/ERK signaling, and results in cell death from pro-apoptotic Death Receptor 5 (DR5) receptor activation [14]. ONC201 has shown to be an agonist of mitochondrial caseinolytic protease (ClpP) [16], which impairs oxidative phosphorylation and leads to cell death and mitochondrial disruption [14]. Additionally, ONC201 is an antagonist of G protein coupled receptor (GPCR) dopamine receptor D2 (DRD2) and D3 (DRD3) [14]. ONC201 is active in various malignancies including H3K27M-mutated glioma and neuroendocrine tumors (NETs) expressing high levels of DRD2 [17]. More recently, a phase II clinical trial study of ONC201 in neuroendocrine tumors with high DRD2 levels, including pheochromocytoma-paraganglioma (PC-PG) and desmoplastic small round cell tumor (DRSCT), showed clinical impact and was well tolerated by patients [18].

The present study investigates the hypothesis that markers of NED, activation of the integrated stress response (ISR) and the TRAIL pathway, as well as the expression of ClpP contribute to neuroendocrine tumor cell death and sensitivity to ONC201/TIC10. By investigating basal NED marker expression in various prostate and SCLC cell lines and determining their sensitivity to ONC201, we aim to better characterize PC and SCLC lines and understand potential molecular targets. We demonstrated that prostate cancer and small cell lung cancer cells are sensitive to ONC201 in both cells with and without high neuroendocrine differentiation characteristics. Our results provide the background for developing ONC201 into clinical studies as therapies for NEPC and NETs.

## Materials and Methods

### Reagents

ONC201 was provided by Oncoceutics/Chimerix, Inc. (Philadelphia, PA, USA) and was stored at -20°C at a concentration of 20 mM.

### Cell culture and growth conditions

Prostate cancer cell lines include PC3, DU145, LNCaP, and 22RV1. Small cell lung cancer cells include H1417 and H1048. All cell lines were acquired from American Type Culture Collection (ATCC). PC3, DU145, 22RV1, LNCaP, and H1417 were cultured in RPMI-1640 medium (Cytiva HyClone SH30027LS) with 10% fetal bovine serum (FBS) and 1% penicillin/streptomycin (P/S). H1048 were cultured in DMEM medium supplemented with 10% FBS and 1% P/S. All cell lines were incubated at 37°C with 5% carbon. Cells were trypsinized with 0.25% trypsin when confluent. Cell line authentication and mycoplasma infection testing were done using PCR testing.

### Cell viability assays

Cells were plated at a density of 3000 cells per well in a 96-well plate and in 100 μL of medium. After 24 hours, 20 μL of ONC201 were added in a 1:2 serial dilution from 20-0 μM and then incubated for 72 hours. 20 μL of CellTiterGlo bioluminescence agent was added to each well and plates were mixed for 3 minutes on a plate shaker to begin cell lysis. Cell viability was collected using a CellTiterGlo assay, and bioluminescence images were collected with the Xenogen IVIS imaging system. Dose response curves and IC50 doses were determined and graphed using GraphPad Prism.

### Colony formation assays

After optimizing experimental conditions of varying cell lines, DU145 and 22RV1 were plated at a density of 200 cells per well of a 12-well plate. PC3 cells were plated at a density of 200 cells per well in a 6-well plate. After 24 hours, cells were treated with a range of doses of ONC201. DU145 and 22RV1 were treated with 1.33, 1.67, and 2 μM of ONC201. PC3 cells were treated with 2,4, and 6 μM of ONC201. After 7 days of incubation, wells were washed with phosphate-buffered saline (PBS), fixed with formalin for 30 minutes, then stained with Coomassie Brilliant blue for 20 minutes. Colonies were counted and percent area was determined using ImageJ.

### Western blot analysis

Neuroendocrine differentiation markers, BRN2/SOX2 transcription factors, and TRAIL pathway proteins in prostate cancer and small cell lung cancer cell lines were examined using a western blot. 500,000-1,000,000 cells were plated and incubated in a 6-well plate for 24 hours for cell adherence. Cells were treated with ONC201 at IC50 drug dose and incubated at different time points (12, 24, 48 hours). After cells were harvested, they were washed with cold PBS and lysed with RIPA buffer with protease inhibitor (Cell Signaling Technology 9806S) and phosphatase inhibitor (PHOSS-RO Roche 4906845001). Protein quantification was done with Pierce BCA Protein Assay (ThermoFisher Scientific, Waltham, MA). Samples were boiled at 95°C for 10 minutes after denaturing sample buffer was added. Cell lysates were electrophoresed through 4-12% SDS-PAGE gels (Invitrogen) and transferred to PVDF membranes. Membranes were blocked in 5% milk in 1X TTBS and incubated overnight at 4°C with primary antibodies in 5% non-fat milk. Primary antibodies were washed and incubated in appropriate HRP-conjugated secondary antibody at a 1:10,000 dilution in 5% non-fat milk (rabbit, ThermoFisher Scientific, or mouse, ThermoFisher Scientific) for two hours. ECL western blotting detection reagent and Syngene imaging were used to measure protein expression levels.

### Transient overexpression

POU3F2-Human_pcDNA3.1(+)-P2A-eGFP plasmid was created from GenScript. Bacteria containing the plasmid from agar stabs were streaked onto LB agar petri dishes with 100 μg/mL ampicillin and grown at 30°C for 24 hours. Single colonies were Midi-prepped (QIAGEN Plasmid Plus Midi Kit, 12943) based on the manufacturer’s protocol. Whole-plasmid sequencing was done to confirm the plasmid sequence (Plasmidsaurus).

### Statistical analysis

Cell viability assays and colony formation assays were analyzed with mean ± SD. ONC201 dose response curves were created with GraphPad. Cell viability assays and western blot analyses were conducted in replicates of three. Colony formation assays were conducted in replicates of nine. Treatment groups for colony formation assays were analyzed with a one-way ANOVA and t-tests with p<0.05 considered statistically significant. GraphPad Prism Version 10.2.2 was used for statistical analysis.

## Results

### ONC201 shows sensitivity and cytotoxicity to prostate and small cell lung cancer cell lines (N=6) at low doses

To understand the sensitivity of TRAIL-inducing ONC201 therapeutic in cell lines, we conducted a cell viability assay in N=6 prostate and small cell lung cancer cells lines. Loss of viability was seen at different drug concentrations, but all cell lines showed decreased viability at low doses of therapeutic. Increased concentrations of ONC201 treated resulted in decreased cell viability in a dose-dependent manner. The most sensitive cell line among the panel was H1417 (IC50 = 1.02 μM) small cell lung cancer and 22RV1 prostate cancer (IC50 = 1.16 μM), followed by H1048 (IC50 = 1.26 μM) and LNCaP (IC50 = 1.31 μM).

**Figure 1.**
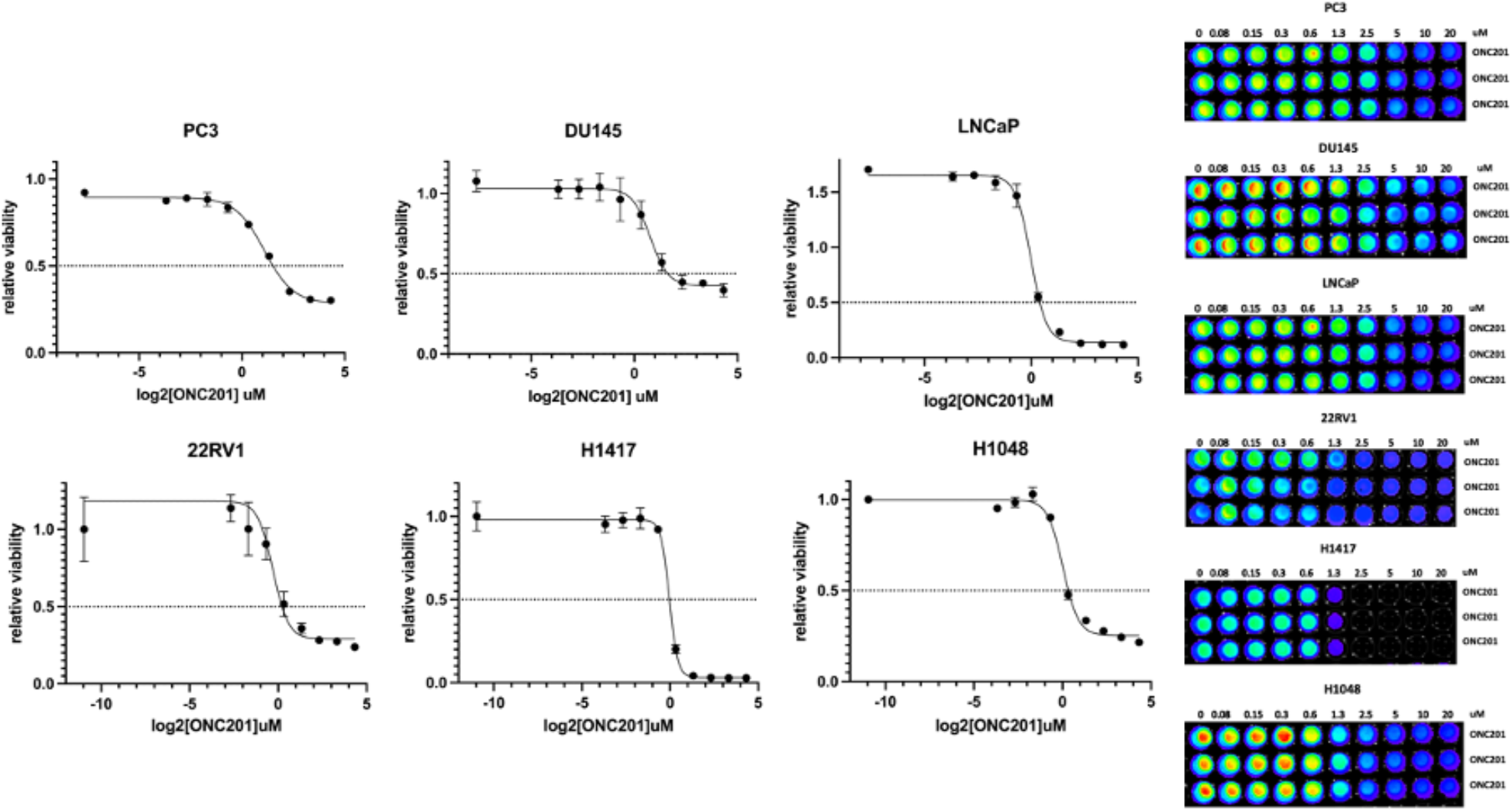
Prostate and small cell lung cancer cell lines IC50 curves after ONC201 treatment for 72 hours. Cell viability was measured through the addition of CellTiterGlo (CTG) reagent. IC50 curves of cell lines presented on the left. Representative images of cell viability on the right.

**Table 1.** IC50 values of cell lines treated with ONC201.

| Cell Line | IC50 value (μM) |
| --- | --- |
| PC3 | 2.87 |
| DU145 | 3.10 |
| LNCaP | 1.31 |
| 22RV1 | 1.16 |
| H1417 | 1.02 |
| H1048 | 1.26 |

### Classification of cell lines on neuroendocrine spectrum model based on basal expression levels of neuroendocrine differentiation markers

To understand the presence and endogenous neuroendocrine markers are present in PC and SCLC, we performed western blots on solid tumor cell lines investigating NED markers SOX2, PGP9.5, and BRN2. In addition, we investigated endogenous levels of TRAIL pathway protein expression levels, specifically ClpX, ClpP, DR5, and cPARP. In PC3, low levels of BRN2 and PGP9.5 was present. In DU145, BRN2 was not expressed, but high levels of PGP9.5 was present. In LNCaP and 22RV1, neither BRN2 nor PGP9.5 was present. In H1417, high levels of SOX2, BRN2, and the highest levels of PGP9.5 compared to all other cell lines were present. In H1048, SOX2 was not present, but BRN2 and PGP9.5 was present, with the highest expression of BRN2 expressed compared to all other cell lines.

**Figure 2.**
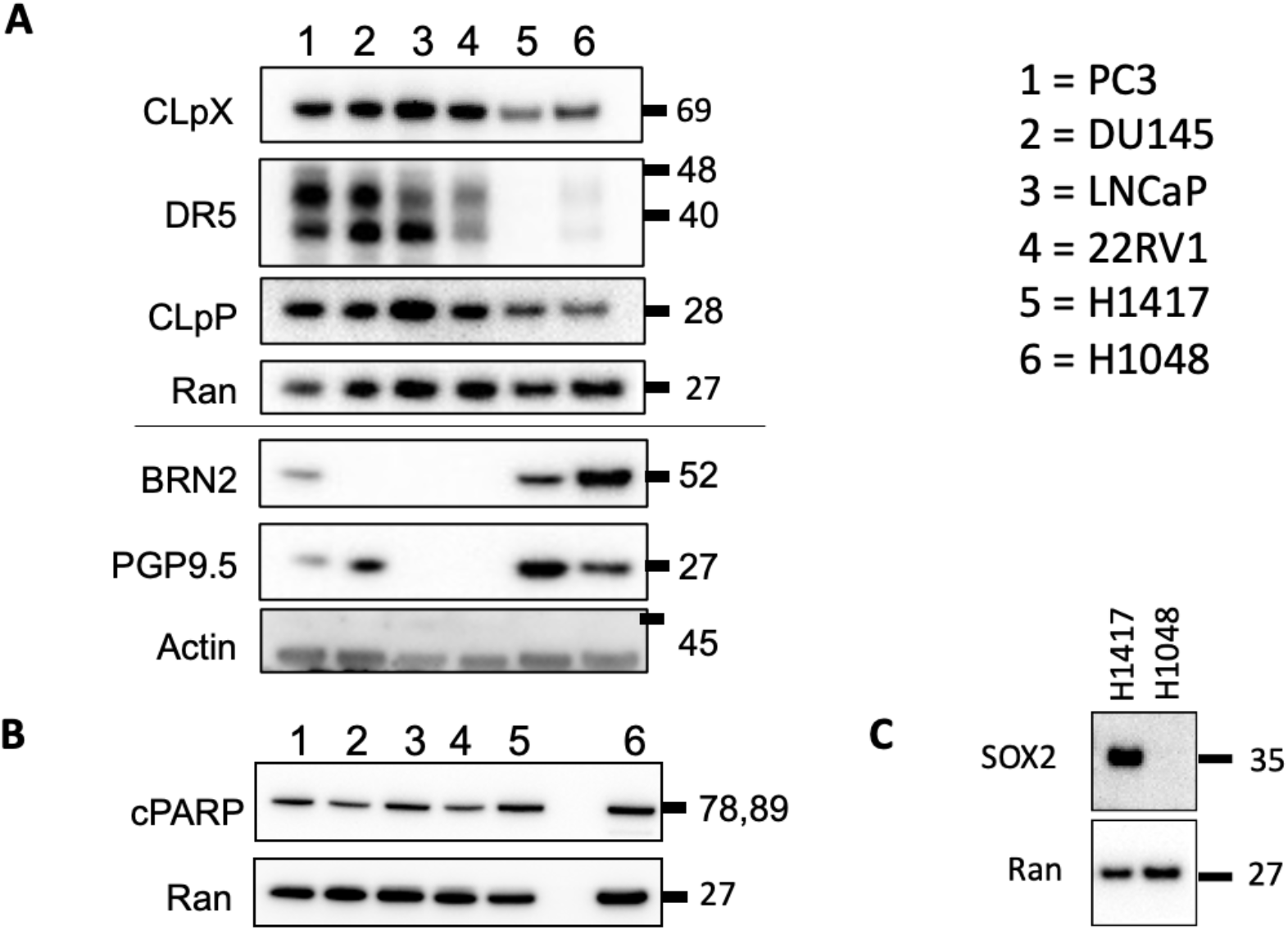
Protein immunoblotting assay showing basal expression levels of neuroendocrine differentiation and TRAIL markers. For all cell lines, all membranes were collected from the same set of lysates and probed with a loading control. Ran and actin was used as a loading control.

**Figure 3.**
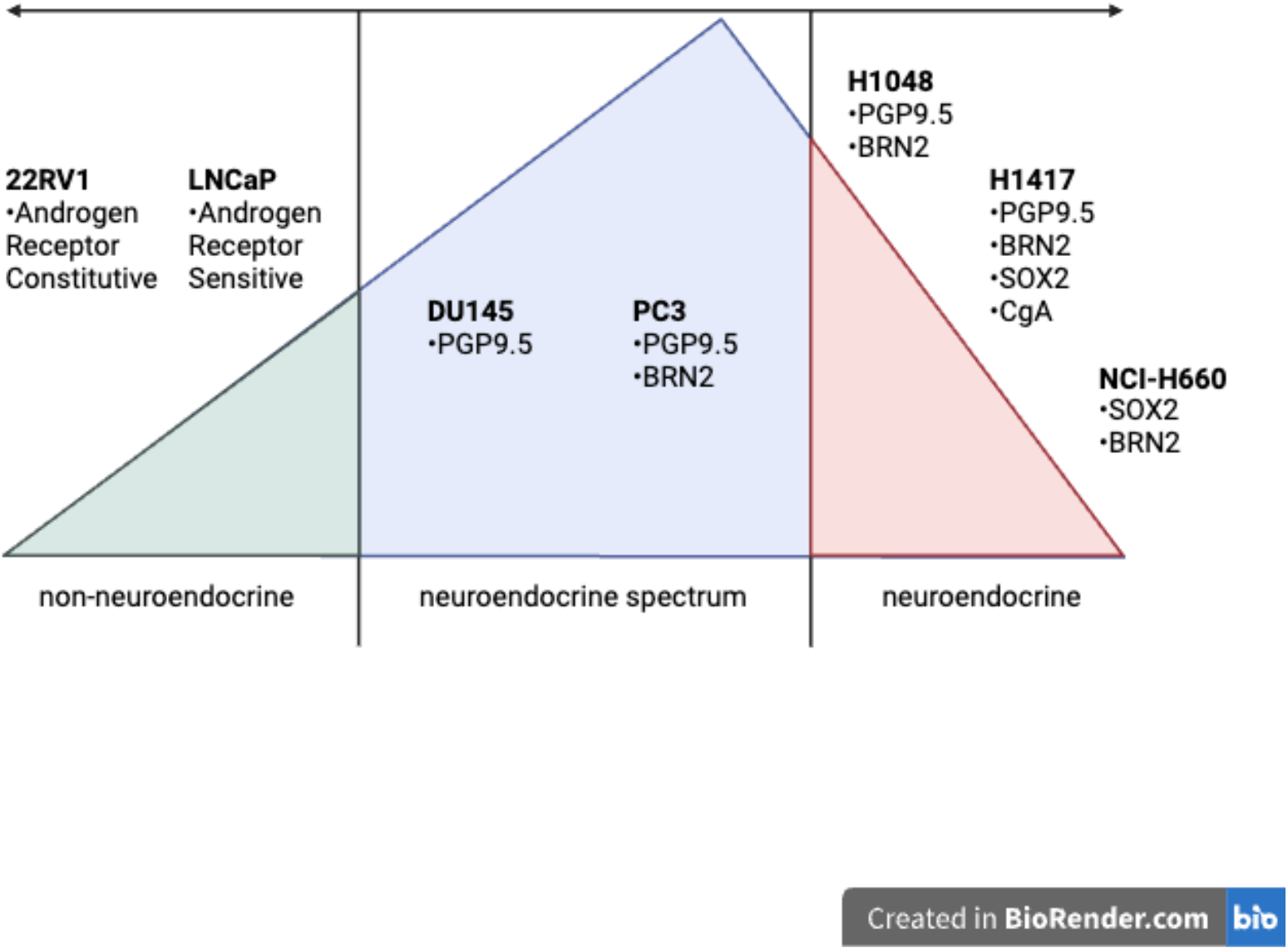
Proposed model of neuroendocrine characterization level of cell lines. Characterization is based on protein expression levels of neuroendocrine differentiation markers of interest and scientific literature.

**Table 2.** Summary of cell lines and their neuroendocrine features.

| NED Marker | PC3 | DU145 | LNCaP | 22RV1 | H1417 | H1048 |
| --- | --- | --- | --- | --- | --- | --- |
| BRN2 | Low levels | ND | ND | ND | Low levels | High levels |
| SOX2 | Low levels | Low levels | Low Levels | Low levels | High levels | ? |
| Chromogranin A (CgA) | ND | ND | ND | ND | High levels | ? |
| PGP9.5/<br>UCLH1 | Low levels | High levels | ND | ND | High levels | Low levels |
| FOXO1 | Low levels | Low levels | High levels | High levels | ? | ? |
| Enolase<br>2/NSE | ND | Low levels | High levels | High levels | ? | ? |
| Androgen<br>Receptor | ND | ND | High levels | ND* | ? | ? |

Protein expression levels were detected through western blotting.
ND: not detected
?: not tested
\*22RV1 expresses AR receptor, but due to the splice site mutation, was not visible in the western blot.

### Early upregulation of ATF4, cPARP, and DR5 and decreased CLpX expression during apoptosis in prostate cancer cell lines at 24-hour ONC201 treatment

We conducted time point western blots to investigate mechanistic pathways resulting in ONC201-induced cell death. In DU145, increase in DR5 expression was observed around 48 hours.Cleaved PARP (cPARP) expression increased at 24 hours, indicating cell death. Chaperone subunit ClpX, which regulates mitochondrial Clp protease, decreased at the 12- and 24-hour time point, suggesting ONC201 induces cell apoptosis through mitochondrial proteolysis. ATF4 increase expression was observed at 12 hours, with significant increase in expression beginning at 24 hours, indicating the activation of the ISR and tumor cell death by ONC201. In 22RV1, an increase in DR5 expression was present around 24 hours and 48 hours. We observed an increase in cPARP expression at 24 hours. ClpX showed slight decrease in expression at 12 hours. ATF4 expression increased at 12 hours, with high expression levels presented at 48 hours.

**Figure 4.**
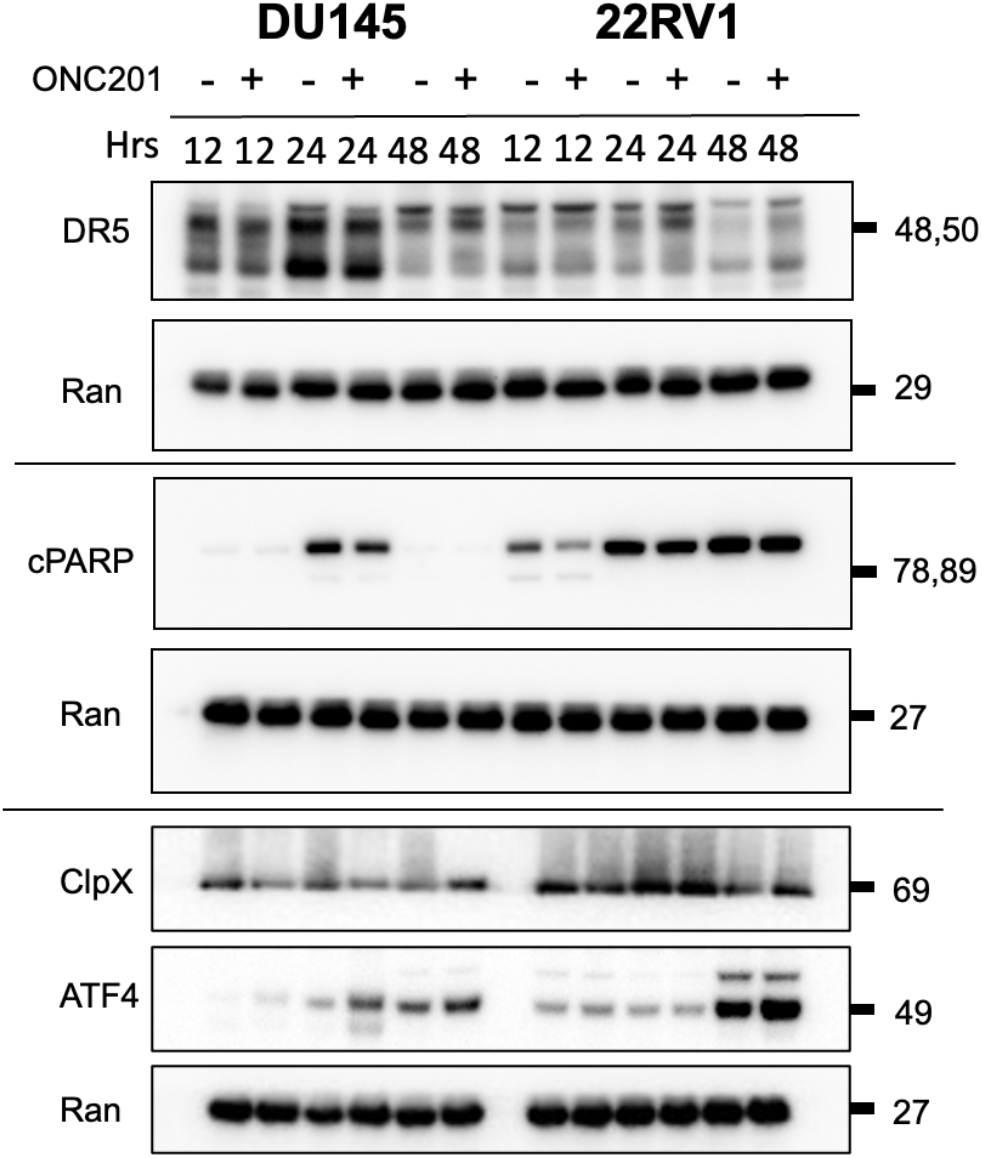
Western blot analysis for extrinsic pathway proteins expression and alteration when treated with IC50 values shown in DU145 and 22RV1 cell lines at 12, 24, and 48 hours. DU145 was treated with IC50 value 3.10 μM ONC201 and 22RV1 was treated with IC50 value 1.16 μM ONC201. For all cell lines, all membranes were collected from the same set of lysates and probed with a loading control.

### ONC201 shows changes in sensitivity and cytotoxicity to prostate cancer cell lines when transiently overexpressed with BRN2

To test if there is a change in sensitivity to ONC201 when BRN2 NE differentiation transcription factor is overexpressed in PC lines, we transiently overexpressed PC3, DU145, 22RV1, and LNCaP cell lines with BRN2 plasmid for 12 hours, treated with ONC201 at various doses, and imaged after 72 hours. 22RV1 had similar sensitivity to ONC201 when BRN2 was overexpressed compared to the control vector but had a slight increase in IC50 value (control vector IC50 = 1.34, BRN2 OE vector IC50 = 1.91). DU145 displayed an increase sensitivity to ONC201 when BRN2 was overexpressed compared to the control vector (control vector IC50 = 4.46, BRN2 OE vector IC50 = 3.85). PC3 and LNCaP showed decrease in sensitivity to ONC201 when BRN2 was overexpressed. Overall, we did not observe a meaningful difference in drug sensitivity.

**Figure 5.**
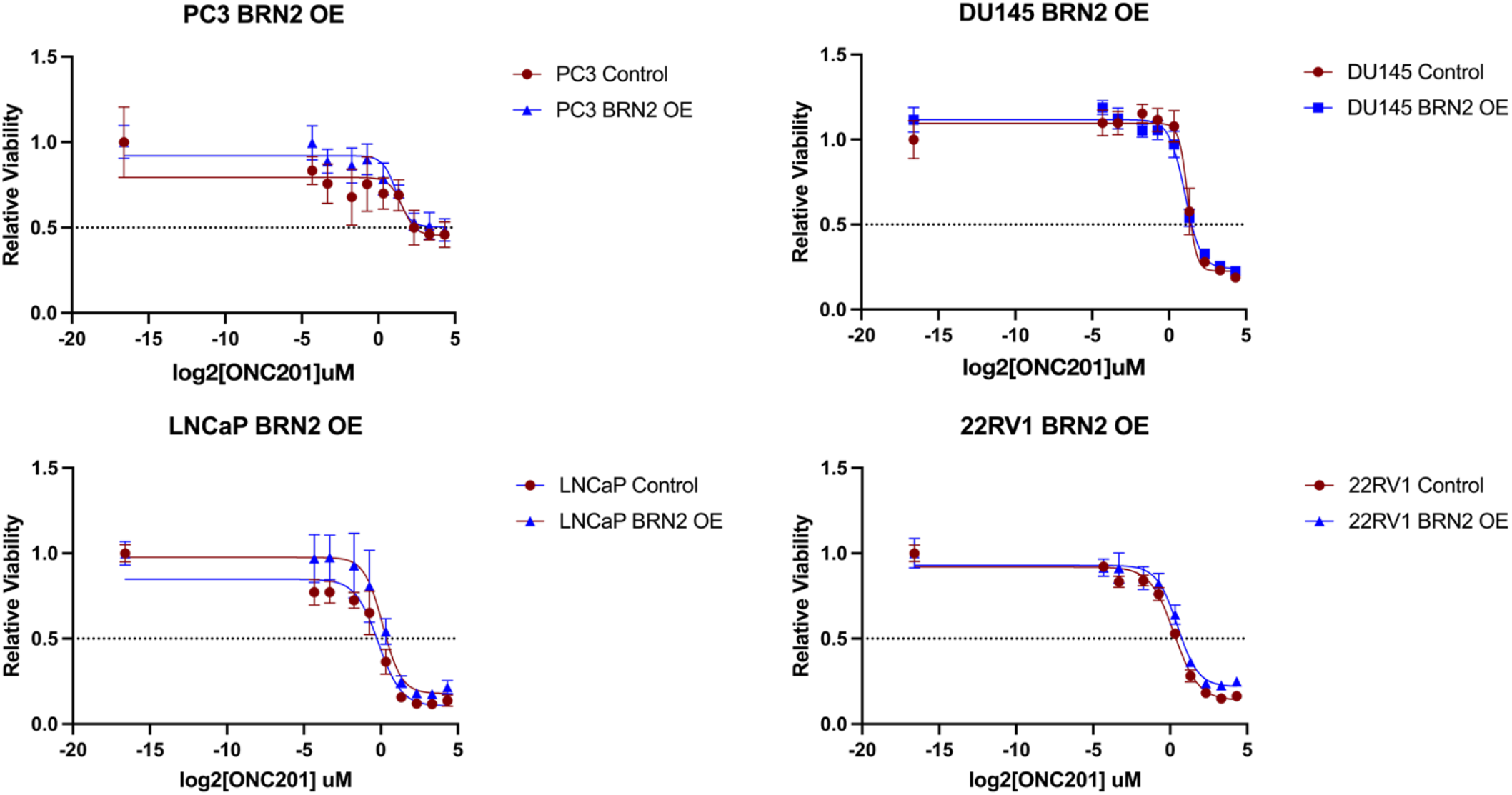
Prostate cancer cell line sensitivity to ONC201 after transient BRN2 overexpression. Cells were transiently overexpressed with 0.8 ug/uL BRN2 plasmid and 0.8 ug/uL pcDNA 3.1 eGFP control plasmid for 48 hours before treated with ONC201 for 72 hours. Cell viability was measured through the addition of CellTiterGlo (CTG) reagent.

**Table 3.** IC50 values of PC lines treated with varying doses of ONC201 with and without transient BRN2 overexpression.

| Cell Line | BRN2 Control<br>IC50 (μM) | BRN2 OE<br>IC50 (μM) |
| --- | --- | --- |
| PC3 | 4.66 | 8.51 |
| DU145 | 4.46 | 3.85 |
| LNCaP | 0.78 | 1.47 |
| 22RV1 | 1.34 | 1.91 |
| H1417 | N/A | N/A |
| H1048 | N/A | N/A |

### Transient overexpression with BRN2 decreases neuroendocrine markers FoxO1 and SOX2 protein expression in prostate cancer cell lines

To further investigate BRN2 and its role in driving neuroendocrine differentiation and regulating neuroendocrine markers expression, we performed western blot analyses in prostate cancer cell lines with transient BRN2 overexpression. We hypothesized that BRN2 overexpression would alter neuroendocrine marker levels. In PC3, we observed a slight increase in FoxO1 levels with BRN2 overexpression, but no change in Enol-2, PGP9.5, or SOX2. Interestingly, for DU145 cell lines, we found a decrease in protein expression levels of FoxO1, Enol-2, and PGP9.5 NED markers when BRN2 was overexpressed. In LNCaP and 22RV1, we observed no change in Fox)1, Enol-2, and PGP9.5 NED markers. However, in LNCaP, we observed a decrease in SOX2 expression. 22RV1 cell lines have no change in SOX2 expression.

**Figure 6.**
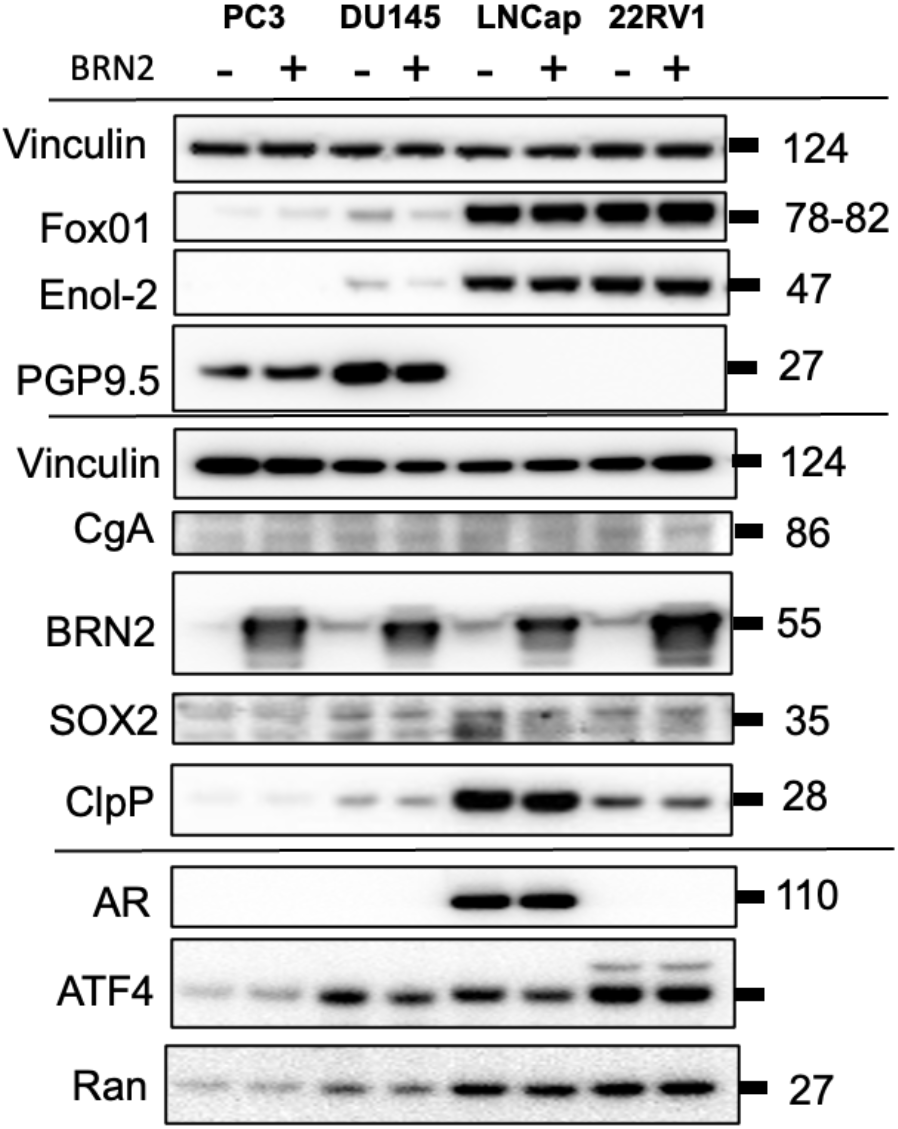
Overexpression of BRN2 is associated with changes in NED marker protein expression at 48 hours. Cells were transiently overexpressed with 0.8 μg/μL BRN2 plasmid and 0.8 μg/μL pcDNA 3.1 eGFP control plasmid for 48 hours. Membranes were probed with a loading control. All membranes were analyzed from one set of lysates.

### BRN2 overexpression with 48-hour ONC201 treatment is associated with a decrease in ClpX expression, increased DR5 expression, and a greater increase in BRN2 expression

For DU145, we observed a decrease in ClpX when ONC201 was added around 48 hours. Additionally, we observed an increase DR5 in both DU145 and LNCaP when ONC201 was added. There was not a significant difference in protein expression levels in PGP9.5 in DU145 when BRN2 was overexpressed. Interestingly, in both DU145 and LNCaP cells that were overexpressed with BRN2, we noticed an increase in BRN2 protein expression levels cells when treated with ONC201.

**Figure 7.**
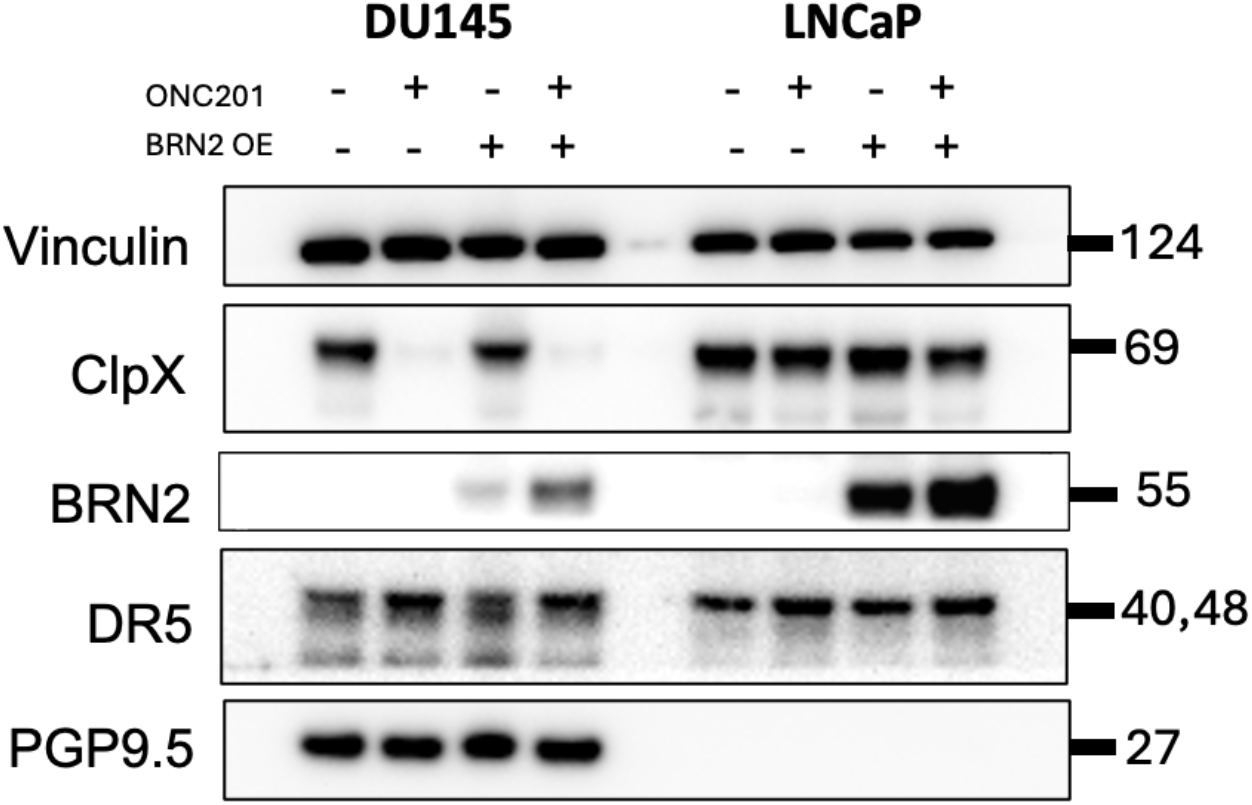
BRN2 overexpression with ONC201 treatment NED markers and TRAIL protein expression in prostate cancer cells. Cells were transfected with 0.8 μg/μL BRN2 plasmid and 0.8 μg/μL pcDNA 3.1 eGFP control plasmid for 48 hours before adding the respective doses of ONC201 for 48 hours. Membrane was probed with a loading control. All membranes were analyzed from one set of lysates.

### ONC201 significantly reduces colony forming ability of prostate cell lines

In addition to cell viability assays, we investigated whether similar sensitivity and response may occur in prostate cancer cell lines using long-term colony formation assays. We observed similar observations in Figure 7 and Table 5 that ONC201 is sensitive to prostate cancer cell lines PC3, DU145, and 22RV1, further supporting cell line sensitivity to ONC201 at low doses. PC3 cells showed significant decrease in colony forming ability at 2.00 μM, with one-way ANOVA tests showing *P* values of <0.0001. DU145 and 22RV1 cells showed significant decrease in colony forming ability at 1.67 μM, with one-way ANOVA tests showing *P* values of <0.0001.

**Figure 8.**
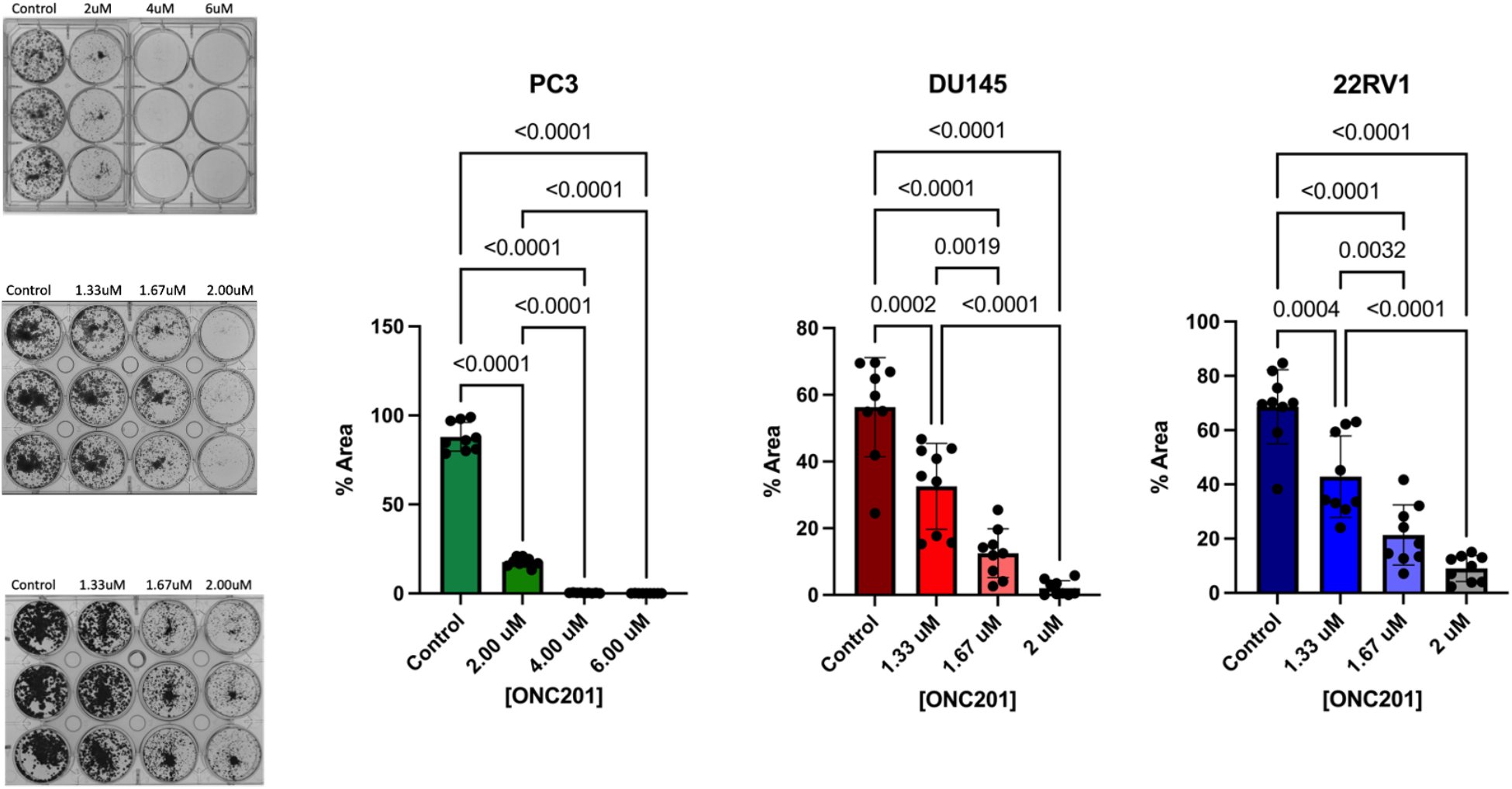
Colony forming ability in prostate cancer cell lines following ONC201 treatment. Images are representative of biological replicates after 7 days (left). Colony formation assay quantification for PC3, DU145, 22RV1 cell lines (right). Colonies are reported as mean ± SD, and treatment groups were compared using one-way ANOVA tests.

**Table 4.** Descriptive statistics showing percent area of PC lines when treated with varying doses of ONC201.

| Cell Line | Control (percent area) | Control to 1.33 | Control to 1.67 | Control to 2 | Control to 4 | Control to 6 |
| --- | --- | --- | --- | --- | --- | --- |
| PC3 | 87.99% ± 8.02 | N/A | N/A | <0.0001 | <0.0001 | <0.0001 |
| DU145 | 56.32% ± 14.87 | 0.0002 | <0.0001 | <0.0001 | N/A | N/A |
| 22RV1 | 68.64% ± 13.68 | 0.0004 | <0.0001 | <0.0001 | N/A | N/A |

## Discussion

Our studies reveal that some prostate cancer cell lines show higher neuroendocrine differentiation characteristics than others, like PC3, DU145, and H1417. However, all cell lines (N=6) are sensitive to ONC201 at low dose ranges, suggesting that ONC201 is efficacious to prostate and small cell lung cancer cells regardless of their NED characteristics. It is important to acknowledge that we are still understanding what characteristics constitute cells being neuroendocrine-like, and more investigation is needed to get a full understanding of what phenotypes occur when cells differentiate. Our western blot analyses show that tumor cell lines possess neuroendocrine differentiation markers of interest.

Our data show that DU145 with transient BRN2 overexpression at 48 hours does not result in greater expression of NEPC markers. However, overexpression slightly increases DU145 sensitivity to ONC201 (**Table 3**) and was evident without an increase in common phenotypical markers used for NEPC assessment (FOX01, ENO2, PGP9.5, CgA) at 48 hours. Our data revealed that protein expression change was not present when PCa had an integral androgen receptor pathway. In 22RV1 and LNCaP, there was no change in markers like Fox01, Enol-2, or PGP9.5. BRN2 overexpression decreased sensitivity of LNCaP, PC3, and 22RV1 to ONC201 (**Table 3**).

A limitation in this study is the lack of *in vivo* work conducted. In addition, our model of BRN2 transient overexpression using plasmids may not be fully representative of the process of neuroendocrine differentiation due to its short time point, and we plan to conduct experiments with stable cell lines with BRN2 and SOX2 overexpressed. More time points should be investigated to see if there are changes in NED marker levels. Another limitation is the number of cell lines that were used in the experiment, and a potential solution in the future is to look at a larger number of cell lines in the future.

### Preliminary siRNA knockdown of transcription factor BRN2 in PC3 cell line indicates successful decrease in protein expression of BRN2 in 5, 10, and 20 pmol siRNA

To perform future BRN2 knockdown experiments, we conducted an optimization experiment on PC3 cells to investigate a concentration of siRNA that would result in significant BRN2 protein expression decrease. We observed decrease in BRN2 protein expression in all concentrations of siBRN2.

**Figure 9.**
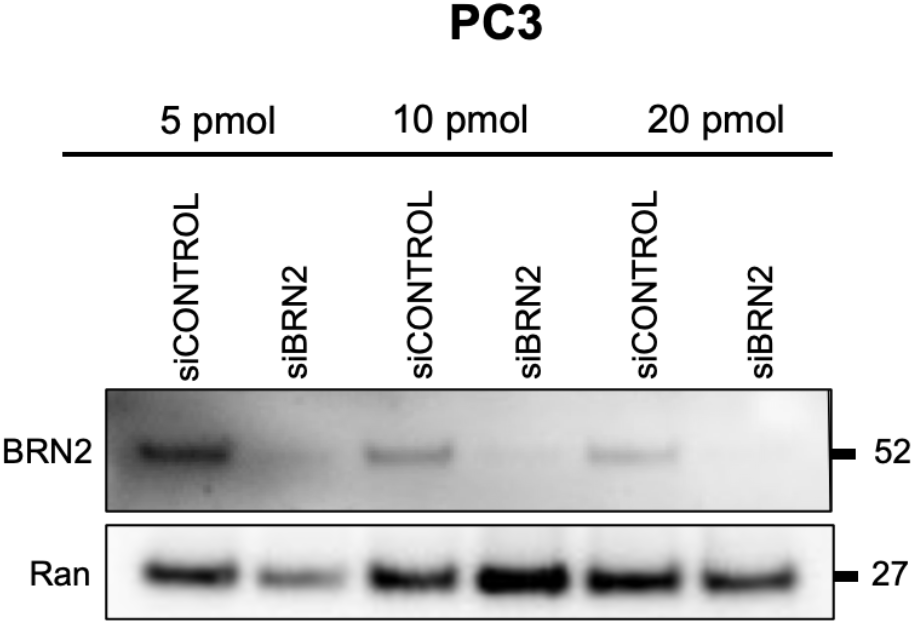
Optimization of BRN2 siRNA concentrations in PC3 prostate cancer cell line. PC3 prostate cancer cells were transfected with varying doses of 5, 10, or 20 pmol of siBRN2 or siCONTROL for 48 hours.

Future directions will evaluate the effects of NE features after overexpression of SOX2 and its sensitivity to ONC201. In addition, future investigations will investigate potential changes of NE features expression levels after siRNA knockdown of BRN2 and SOX2 in prostate and SCLC lines. In addition, we plan to knockdown multiple determinants of ONC201/TIC10 response to evaluate the impact on drug sensitivity. Immunostaining at various time points in PC cells will be done to further understand ONC201 sensitivity. Finally, we will investigate *in vivo* experiments. Overall, our data show evidence that human solid tumor prostate cell lines show sensitivity to ONC201, with varying levels of NED features based on protein expression. It is important that neuroendocrine tumors are sensitive to ONC201, but equally important to recognize that prostate cancer and small cell lung cancer with and without strong neuroendocrine features are also sensitive to ONC201. Results from our work provide in clinical trials of prostate cancer testing imipridones, our studies would suggest that non-neuroendocrine patients would be eligible and should be includes in clinical trials.

## Acknowledgments

W.S.E-D. is an American Cancer Society Research Professor and is supported by the Mencoff Family University Professorship at Brown University. This work was supported by an NIH grant (CA173453) to W.S.E-D. The contents of this manuscript are solely the responsibility of the authors and do not necessarily represent the official views of the National Cancer Institute, the National Institutes of Health, or the American Cancer Society.

